# Crosstalk Between ALPK1 and STING: A Synergistic Axis in Innate Immune Activation and Human Inflammatory Disease

**DOI:** 10.1101/2025.06.30.662363

**Authors:** Chong-Shan Shi, Christina Kozycki, Ning-Na Huang, Il-Young Hwang, Dima A. Hammoud, Daniel L. Kastner, John H. Kehrl

**Affiliations:** B-cell Molecular Immunology Section, Laboratory of Immunoregulation and Infectious Diseases, National Institute of Allergy and Infectious Diseases, National Institutes of Health, Bethesda, Maryland 20892; Innate Immune Activation Unit, Laboratory of Clinical Immunology and Microbiology, National Institute of Allergy and Infectious Diseases, Bethesda, Maryland 20892; Inflammatory Disease Section, National Human Genome Research Institute, National Institutes of Health, Bethesda, Maryland, 20892; Radiology and Imaging Sciences, National Institutes of Health Clinical Center, Bethesda, Maryland 20892

**Keywords:** ALPK1, STING, Interferon, Inflammasome, Autoinflammatory, TIFA

## Abstract

Alpha kinase 1 (ALPK1) is a cytosolic sensor of microbial sugar metabolites that activates NF-κB signaling through phosphorylation of the adaptor protein TIFA. Although canonically linked to NF-κB, individuals with gain-of-function ALPK1 mutations also show features of interferon-driven inflammation. Here, we show that ALPK1 activation enhances multiple outputs of the stimulator of interferon genes (STING) pathway, including both canonical and noncanonical responses such as STING proton channel–dependent LC3B lipidation and NLRP3 inflammasome activation. Furthermore, ALPK1 signaling activates eIF2α, an effector of the integrated stress response. Conversely, STING activation increases ALPK1 protein expression and triggers TIFA-Threonine 9 phosphorylation. Clinically, individuals with ALPK1-mediated disease exhibit premature intracranial mineralization and elevated cerebrospinal fluid neopterin, both associated with dysregulated interferon signaling. These findings support a model of bidirectional signaling between ALPK1 and STING, in which microbial and nucleic acid sensing pathways can amplify one another. This crosstalk provides a mechanistic framework for understanding innate immune signaling relevant to both homeostasis and disease.

## Introduction

The innate immune system coordinates host defense through a network of pattern recognition receptors (PRRs) that detect conserved microbial products and endogenous danger signals (Carpenter and O’Neill, 2024). Although often studied in isolation, these pathways do not function independently. Emerging evidence suggests that crosstalk between PRRs is a fundamental property of immune signaling - enabling integration of diverse cues, tailoring of inflammatory responses to tissue context, and coordination of downstream metabolic and regenerative programs. Understanding these interactions is essential not only for elucidating immune signaling architecture but also for identifying new therapeutic targets in complex inflammatory and degenerative diseases.

One example highlighting the potential relevance of PRR crosstalk is alpha-protein kinase 1 (ALPK1), a vertebrate-restricted PRR that has only recently been characterized (Tang et al., 2024; Zhou et al., 2018). ALPK1 was initially identified as a cytosolic sensor of bacterial metabolites involved in lipopolysaccharide (LPS) biosynthesis including adenosine diphosphate-(ADP)-heptose. Upon ligand binding, ALPK1 phosphorylates TIFA (tumor necrosis factor receptor-associated factor-interacting protein with a forkhead-associated domain), triggering TIFA oligomerization, recruitment of TRAF6 (TNF receptor-associated factor 6), and activation of the transcription factor nuclear factor-κB (NF-κB) (Milivojevic et al., 2017; Zhou et al., 2018; Zimmermann et al., 2017). Although originally recognized as a bacterial sugar sensor, it has subsequently been shown that organisms across diverse biological kingdoms—including bacteria, archaea, eukaryotes, and viruses—encode enzymes capable of producing heptose metabolites that activate ALPK1 (Tang et al., 2024). These findings broaden the physiological relevance of ALPK1 and underscore its potential to shape immune responses across a wide range of infectious and inflammatory contexts.

While ALPK1 is canonically linked to NF-κB signaling, the full scope of its downstream signaling remains incompletely characterized. Insights from human genetics suggest that ALPK1 activation may engage additional immune pathways beyond canonical NF-κB signaling. ROSAH syndrome (retinal dystrophy, optic nerve edema, splenomegaly, anhidrosis, and headache), a monogenic autoinflammatory disease caused by gain-of-function mutations in ALPK1, features both classic NF-κB-associated phenotypes such as anhidrotic ectodermal dysplasia and atypical features not typically associated with NF-κB dysfunction (Kozycki et al., 2022). For instance, premature basal ganglia mineralization, a radiographic hallmark of type I interferonopathies such as Aicardi-Goutières syndrome and stimulator of interferon genes (STING)-associated vasculopathy with onset in infancy (SAVI) and congenital infection with viruses like cytomegalovirus (CMV) (Alarcon et al., 2013; Fremond et al., 2021; La Piana et al., 2016) has been observed in several individuals with ROSAH. Because ROSAH syndrome results from constitutive ALPK1 activation, it uniquely provides causal insights into the consequences of chronic ALPK1 signaling in humans and reveals previously unrecognized interactions with other innate pathways, including the STING axis.

STING is a cytosolic adaptor best known for sensing foreign or damaged host DNA through cyclic GMP-AMP synthase (cGAS). Upon detecting cytosolic DNA, cGAS catalyzes the synthesis of cyclic GMP-AMP (2′3′-cGAMP), which binds and activates STING. Activated STING translocates from the endoplasmic reticulum (ER) to the Golgi apparatus where it facilitates activation of TANK-binding kinase 1 (TBK1) and interferon regulatory factor 3 (IRF3), inducing transcription of type I interferons and other pro-inflammatory cytokines (Dvorkin et al., 2024). Upon translocation to the Golgi, STING also functions as a proton channel, and the resulting cytosolic acidification promotes light chain 3 (LC3) lipidation and NLRP3 inflammasome activation (Liu et al., 2023). To resolve the response, STING is trafficked from the Golgi to lysosomes for degradation, a tightly regulated process essential for limiting the inflammatory response.

Here, we test the hypothesis that ALPK1 and STING engage in functional crosstalk. Drawing on both clinical observations from an expanded ROSAH cohort and mechanistic studies in cellular models, we demonstrate bidirectional amplification between ALPK1 and STING signaling. We show that ALPK1 activation enhances STING activation by stabilizing STING protein and amplifies the STING-mediated interferon response, increases LC3 lipidation, enhances NLRP3 inflammasome activation, and augments the integrated stress response. Conversely, STING activation increases ALPK1 expression and triggers phosphorylation of its adaptor protein TIFA. Together, these findings reveal a previously unrecognized mechanism by which microbial sensing via ALPK1 and nucleic acid sensing via STING may converge to regulate inflammation and shape the immune response.

## Results

### ROSAH patient evaluation suggests dysregulated interferon production

Forty-eight individuals with ROSAH were included in this cohort study (**Table 1**). Of these, 46 individuals from 18 unrelated families were genetically confirmed to be heterozygous for the ALPK1 T237M variant. One individual declined genetic testing but is the child of a genetically confirmed individual with the T237M variant and has optic nerve edema, anhidrosis and splenomegaly and was included in the cohort. Additionally, one individual has the Y254C variant.

**Table 1.**
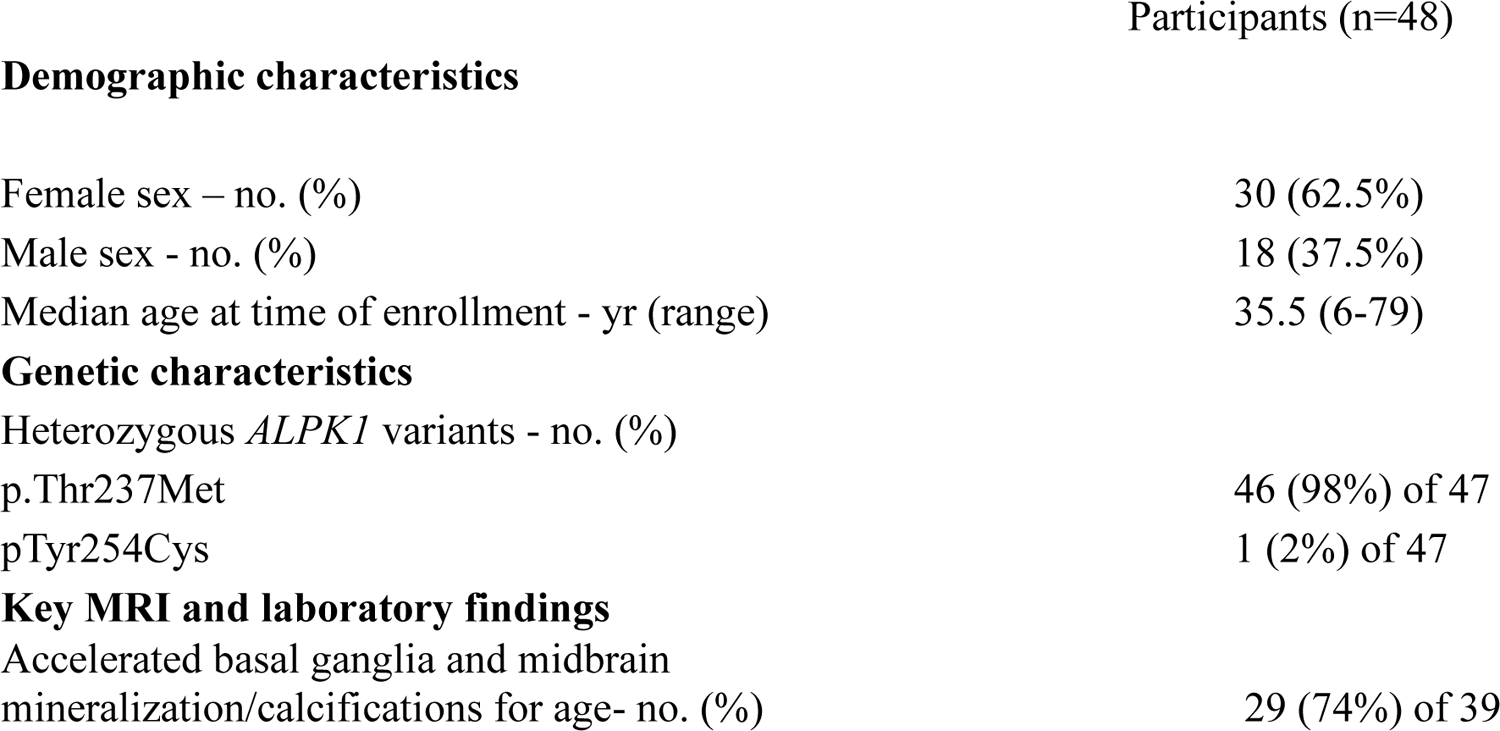
Demographic and Clinical Characteristics of Participants with ROSAH.

**Demographic characteristics**
|  |  |
| --- | --- |
| Female sex – no. (%) | 30 (62.5%) |
| Male sex - no. (%) | 18 (37.5%) |
| Median age at time of enrollment - yr (range) | 35.5 (6-79) |
| **Genetic characteristics** |  |
| Heterozygous *ALPK1* variants - no. (%) |  |
| p.Thr237Met | 46 (98%) of 47 |
| p.Tyr254Cys | 1 (2%) of 47 |
| **Key MRI and laboratory findings** |  |
| Accelerated basal ganglia and midbrain mineralization/calcifications for age- no. (%) | 29 (74%) of 39 |

Among the cohort, 39 individuals had an MRI brain scan available for review. Of these, 29 (74%) exhibited accelerated mineralization/calcification of the basal ganglia, red nuclei and substantia nigra, when compared to controls. A representative montage of the observed abnormalities is shown, in comparison to similar-age healthy control subjects (**Fig. 1A-D**). While basal ganglia mineralization in interferonopathies can present with spasticity and dystonia, individuals with ROSAH—like reports in SAVI—remain clinically asymptomatic despite radiographic findings.

**Fig. 1.**
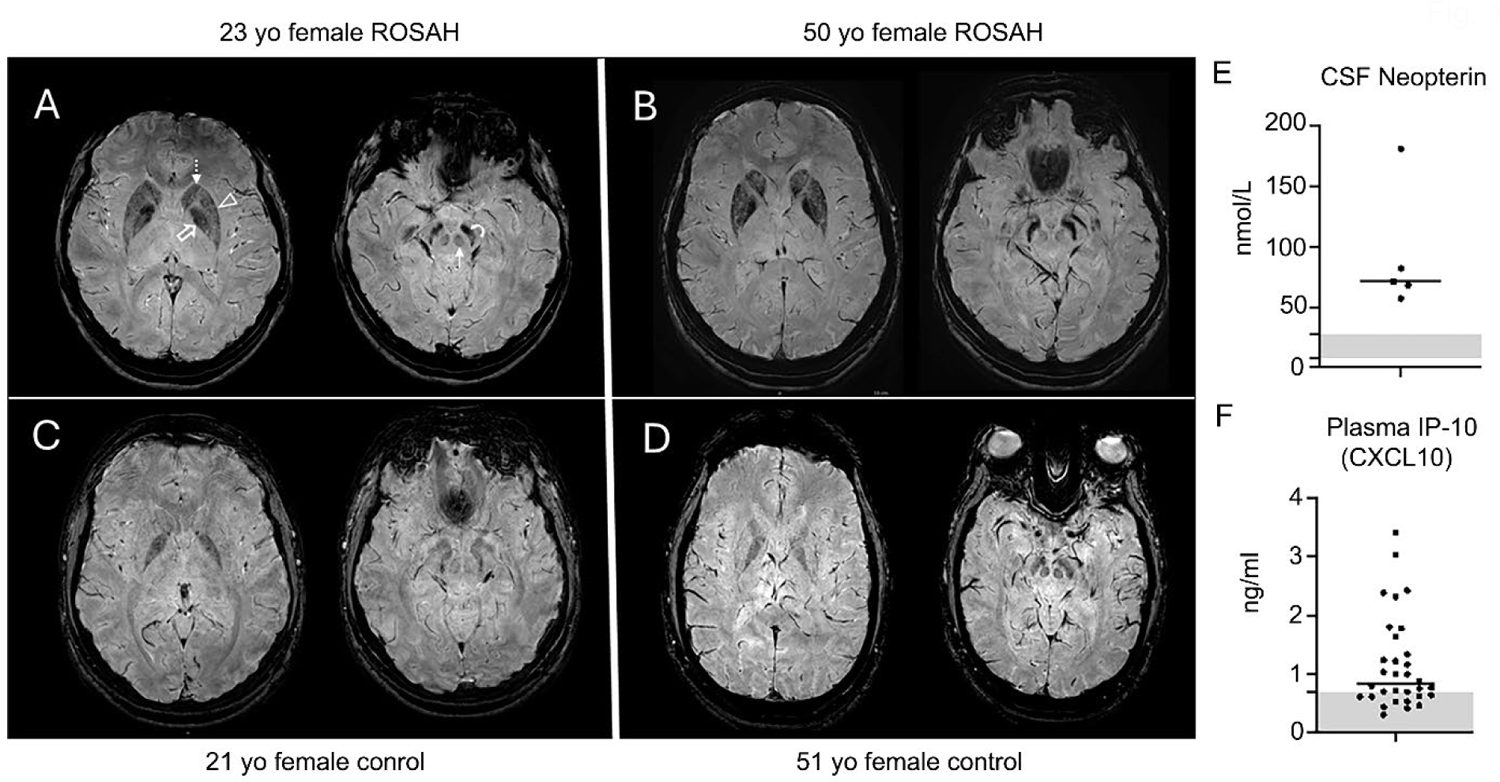
Assessment of features associated with monogenic interferonopathies in individuals with ALPK1 gain-of-function mutations. **(A-D)** Susceptibility weighted images obtained on a 3T MRI scanner at the level of the basal ganglia and midbrain in (A) 23-year-old female and (B) 50-year-old female with ROSAH compared to (C) 21-year-old and (D) 51-year-old healthy control females. There is marked decreased signal intensity in the basal ganglia (caudate (dashed arrow), putamen (arrowhead) and globus pallidus (open arrow) and brainstem structures (red nuclei (solid arrow) and substantia nigra (curved arrow)) of ROSAH patients reflecting accelerated mineralization/calcification. This is more than expected compared to unaffected controls of similar ages. **(E)** CSF neopterin levels in five individuals with ROSAH. (**D**) Plasma CXCL10 levels were elevated in 14 of 20 individuals. The shaded grey areas represent the reference value range.

Cerebrospinal fluid (CSF) neopterin levels were assessed in five individuals, all of whom demonstrated elevated values (**Fig. 1E**). Neopterin, a byproduct of GTP metabolism in interferon-γ stimulated macrophages and monocytes, serves as a key marker of immune system activation, particularly in the context of inflammation in the central nervous system (CNS). Plasma CXCL10 levels were measured in 20 individuals and found to be elevated in 14 (70%) (**Fig 1F**). Seven patients had whole blood RNA analyzed for expression of interferon-induced genes and increased expression was not observed, however, none of the patients had an elevated C-reactive protein at the time of sample collection. Together these data suggests that patients with ROSAH syndrome have dysregulation of interferon signaling, particularly within the CNS compartment.

### ALPK1 activation enhances IFN-β expression via a STING dependent pathway

To evaluate how ALPK1 and STING signaling pathways might interact, we used three cell line HEK 293T, HeLa, and THP-1 cells. We chose HEK 293T cells as they lack cGAS and STING expression (Zhang et al., 2014), but express low levels of ALPK1, HeLa cells as they express very little STING or ALPK1, and differentiated THP-1 cells, a monocyte derived cells line that expresses both ALPK1 and STING (**Fig. 2A**). Initially, we verified that the Y254C and T273M gain-of-function ALPK1 mutant proteins augmented NF-κB transcriptional activity in HEK 293T cells by co-expressing a NF-κB reporter construct along with either wild type ALPK1, or the gain-of-function mutants. As expected, both ALPK1 Y254C and ALPK1 T237M strongly enhanced luciferase activity driven by a NF-κB-responsive promoter (**Fig. 2B**). To check the importance of ALPK1 kinase activity, we expressed a kinase dead version of ALPK1(Zhou et al., 2018), ALPK1 Y254C, or ALPK1 T273M along with the NF-κB reporter. The data showed that the kinase activity of ALPK1 was critical for inducing NF-κB activity (**Fig. 2B**). Next, we expressed an IFN-β reporter construct along with ALPK1 in the presence or absence of ADP-heptose, ALPK1 Y254C, or ALPK1 T237M in HEK 293T cells. In contrast, to the NF-κB reporter ALPK1 and its variants had little effect on the IFN-β reporter gene activity (**Fig. 2C**). Next, we co-expressed STING along with the ALPK1 expression vectors in a similar experiment. Expressing STING alone had an insignificant effect on IFN-β promoter activity, the addition of the ALPK1 construct even in the presence of ADP-heptose had a minimal effect, however, both ALPK1 gain-of-function mutants potently enhanced IFN-β promoter activity in the presence of STING (**Fig. 2C**). Disabling the kinase activity of the ALPK1 gain of function mutants completely inhibited the IFN-β promoter activity previously observed (**Fig. 2D**). To address whether ALPK1-induced NF-κB activation is needed for the activation of the IFN-β promoter construct in HEK 293T cells expressing STING, we used CRISPR to delete expression of the NF-κB family member p65 in HEK 293T cells. As expected, the p65 knock-out severely reduced the NF-κB reporter gene activity triggered by the ALPK1 gain-of-function mutants (**Fig. 2E**). Likewise, the loss of p65 severely impaired the ALPK1 gain-of-function mutants IFN-β activation (**Fig. 2F**). Finally, we transfected STING, either ALPK1 Y254C or ALPK1T237M, stimulated the cells with low concentration of the STING agonist diABZI, and assessed IFN-β promoter activity. diABZI is structurally optimized to fit into the STING binding pocket as do natural cyclic dinucleotides like cGAMP (Ramanjulu et al., 2018). We found that together the ALPK1 gain-of-function mutants and diABZI potently enhanced the activity of the IFN-β promoter (**Fig. 2G**). The above data indicate that the induction of IFN-β promoter activity by the ALPK1 gain-of-function mutants depends on the presence of STING and the induction of NF-κB target genes and can amplify a suboptimal concentration of a direct STING agonist.

**Fig. 2.**
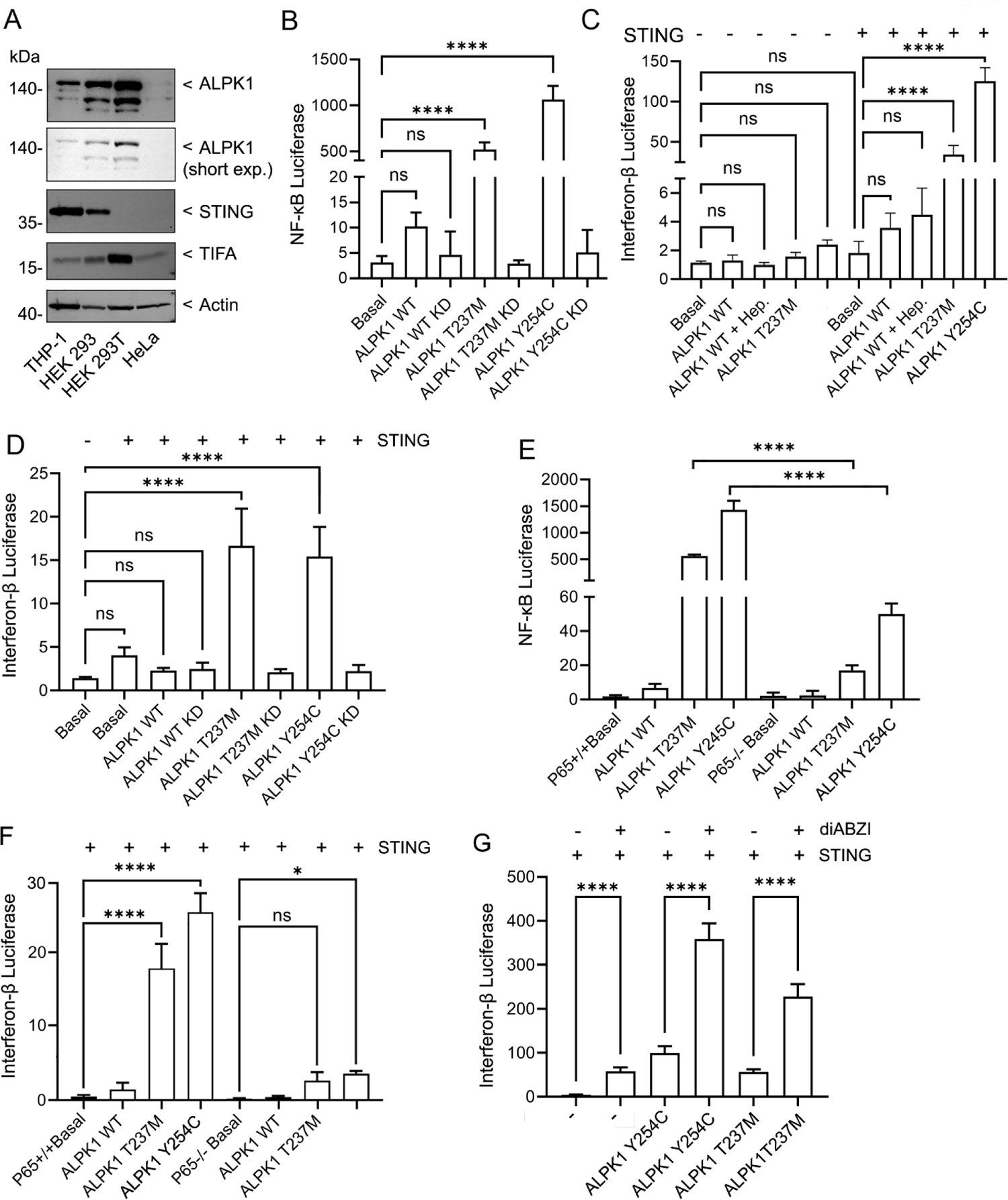
ALPK1 gain-of-function mutations enhanced IFN-β promoter activity required the presence of STING and p65 NF-κB. (**A**) Immunoblots examining ALPK1, STING, and TIFA in cell lysates prepared from 4 different cell lines. Actin immunoblot to assess lysate loading. (**B-G**) HEK 293T cells were transiently transfected with a NF-κB or IFN-β luciferase reporter along with an internal pRL Renilla luciferase control reporter and the indicated plasmids. The results are shown as fold changes. (**B**) The effect of expressing ALPK1 wild type, gain of function mutants T237M and Y254C, and their corresponding kinase-dead forms on NF-κB luciferase reporter activity (fold changes shown). (**C**) The effect of expressing STING on ALPK1, ALPK1 Y254C, and ALPK1 T237M induction of IFN-β promoter activity. ADP-heptose was used at 1.5 µg/ml overnight. (**D**) Comparison of ALPK1, ALPK1 Y254C, ALPK1 T237M and their respective kinase deficient forms on IFN-β luciferase reporter activity when co-expressed with STING. (**E** and **F**) Comparison of HEK 293T cells deficient in NF-κB p65, or not, on ALPK1, ALPK1 Y254C, and ALPK1 T237M induced NF-κB reporter or IFN-β reporter activity. STING co-expressed in part F. (**G**) Induction of IFN-β reporter by a STING agonist is amplified by co-expressing STING with ALPK1 Y254C or ALPK1 T237M. ****, p<0.0001; p>0.05; NS-non-significant. Mean =+/- S.E.M. of n=6 biological replicates.

### Intracellular localization of STING and ALPK1

To characterize the intracellular localization of ALPK1 and ALPK1 Y254C and to determine whether their expression altered that of STING we first used HeLa cells. We expressed ALPK1 or ALPK1 Y254C tagged at their c-termini with the green fluorescent protein Emerald (ALPK1-Em) in the presence or absence of STING tagged at its c-terminus with the red fluorescent protein mRuby (STING-mRuby). Live cell imaging revealed that ALPK1 and ALPK1 Y254C largely distributed homogenously within the cytosol and did not appreciably affect the distribution of STING, which localized in an ER-like pattern (**Fig. 3A**). Second, to assess whether the acute exposure of cells to ADP-heptose altered the distribution of ALPK1 or STING we co-transfected HeLa cells with ALPK1-EM and STING-mRuby and added ADP-Heptose to the cells during live cell imaging. ADP-Heptose addition did not change the localization or distribution of ALPK1 or STING, however, a gradual increase in STING expression occurred after adding it. The expression of STING remained stable in those cells that did not co-express ALPK1 but rose in those cells that expressed both proteins. In contrast the expression of ALPK1 remained stable following addition of ADP-Heptose (**Fig. 3B-D, supplementary video 1 & 2**). In an occasional cell we saw a gradual co-localization of STING and ALPK1 into large cytoplasmic inclusions (**Fig. 3E, supplementary video 3**). This usually preceded loss of cell viability. These results suggest that activation of ALPK1 with ADP-Heptose or constitutively active ALPK1 increases STING expression or stability without causing a detectable change in its intracellular localization. A loss of cell viability was often accompanied by the accumulation of STING and ALPK1 in cytosolic inclusions. As most of the STING/ALPK1 signaling assays used HEK 293T cells we also imaged STING-mRuby along with ALPK1 Y254C-Em in those cells. To activate the STING signaling pathway we treated the cells with diABZI. We found that in some cells diABZI causes a re-localization of STING without affecting ALKP1. But in others the treatment altered the intracellular localizations of both STING and ALPK1 leading to their juxta nuclear colocalization (**Fig. 3F, supplementary video 4**).

**Fig. 3.**
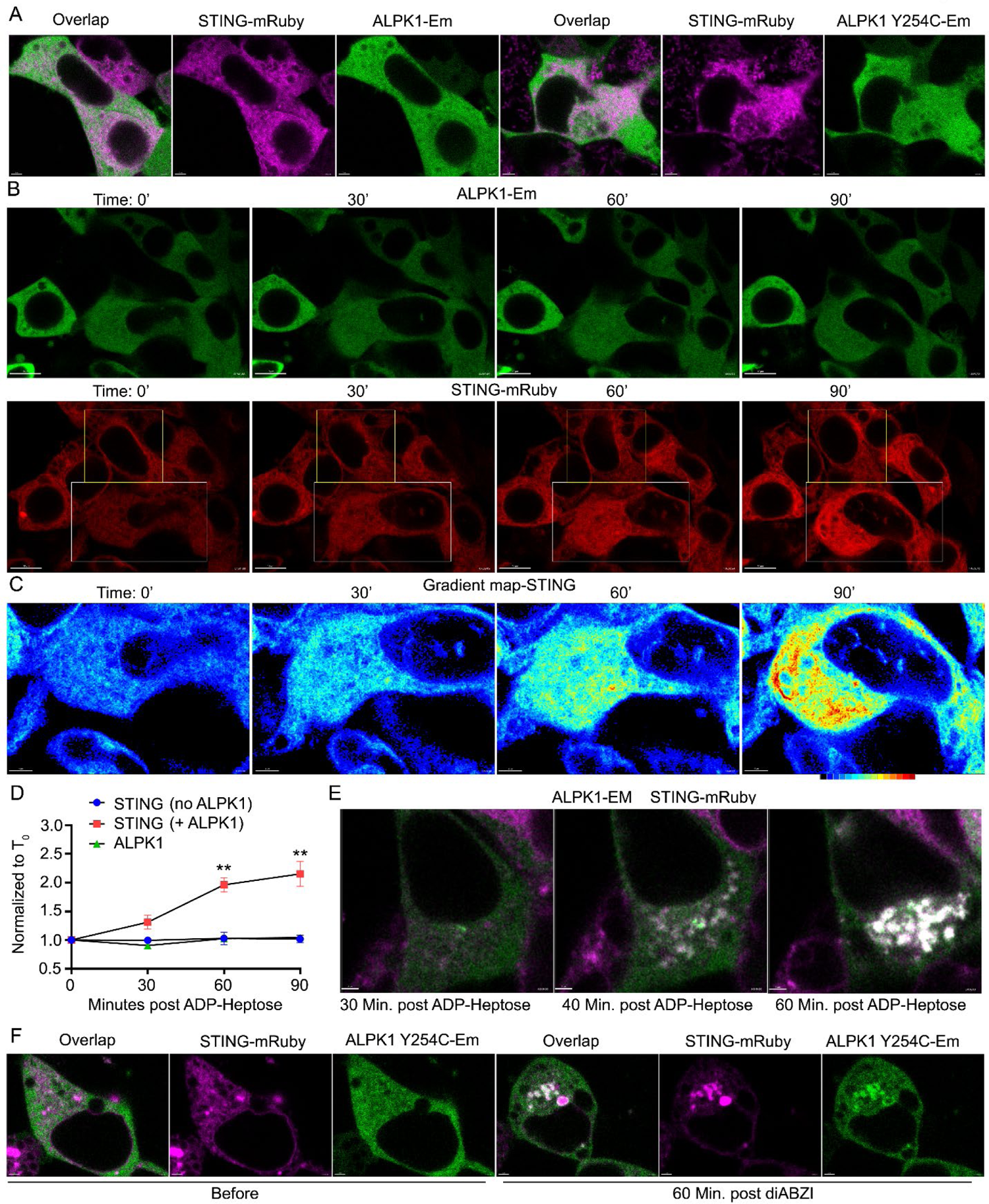
Intracellular localization of ALPK1, ALPK1 Y254C, and STING. HeLa cells (**A-E**) or HEK293T cells (**F**) cells were transiently transfected with STING in the presence of ALPK1 or ALPK1 Y254C and analyzed by confocal microscopy. (**A**) Representative single slice images showing the localization of ALPK1-Em, ALPK1Y254C-Em, and STING-mRuby. STING-mRuby is shown in pink and the ALPK1 constructs in green along with their overlap. Scale bars-3 microns. (**B**) Sequential images showing localization of ALPK1-Em and STING-mRuby before and 30, 60, or 90 minutes after addition of 1.5 µg/ml ADP-heptose. ALPK1-Em (green, above) and STING mRuby (bottom, red). Scale bars-10 microns. (**C**) Sequential zoomed gradient mapped images of STING expression from part B. Gradient shown below 90’ image. Scale bars-4 microns. (**D**) Fluorescent intensity before and 30, 60, or 90 minutes after addition of 1.5 µg/ml ADP-heptose of ALPK1-Em and STING-mRuby. Cells chosen from fields containing double positive and single positive cells post ADP-heptose. Fluorescent intensities normalized to image prior to addition of ADP-heptose. Results from analysis of 15 cells for each condition from two separate transfections. Mean +/- S.E.M. **, p<.005. (**E**) Confocal images of occasional cell showing cytosolic inclusions with strong co-expression of ALPK1 and STING. Overlap images of ALPK1-Em (green) and STING-mRuby (pink) shown at various time points following addition of ADP-heptose (1.5 µg/ml). Scale bar-2 microns. (**F**) Image of cell showing localization of STING with ALPK1 Y254C following addition of diABZI. Cell expressing ALPK1-Y254C Em (green) and STING-mRuby (Pink) and stimulated with diABZI (1 µM). Images before and 1 hour post stimulation. Scale bar-2 microns.

### ALPK1 primes cells to facilitate STING signaling

Following activation STING oligomers translocate from the ER to the Golgi, where they recruit TBK1 promoting TBK1 autophosphorylation and activation (Jeltema et al., 2023). Activated TBK1 phosphorylates STING at position S366 and IRF3 at position S386 (Tanaka and Chen, 2012; Zhang et al., 2019). This leads to IRF3 dimerization and its nuclear translocation (Jeltema et al., 2023). To check whether the gain-of-function mutants affected STING oligomerization we immunoblotted STING in cell lysates that co-expressed STING with wild type ALPK1 or either of the ALPK1 mutant proteins. We found that ALPK1 had no effect, while either gain-of-function version increased the detection of STING oligomers. In addition, STING expression level increased when co-expressed with ALPK1 Y254C or ALPK1 T273M (**Fig. 4A**). Next, we examined the levels of pS366 STING, pS172 TBK1 (Larabi et al., 2013), and pS386 IRF3 (Servant et al., 2003) following expression of ALPK1, plus or minus ADP-heptose; or its gain-of-function mutants. Agonist activated ALPK1 and the two gain-of-function mutants increased the phosphorylation of STING, TBK, and IRF3 on the residues noted above (**Fig. 4B**). Having relied on transfected proteins we examined whether agonist activated ALPK1 facilitated endogenous STING activation in THP-1 cells, a human monocyte/macrophage-like cell line. We primed the cells by treating with phorbol 12-myristate 13-acetate (PMA), which causes THP-1 cells to adopt a macrophage-like morphology and stimulated the cells with the STING agonist diABZI in the presence or absence of the ALPK1 agonist ADP-Heptose. We monitored the phosphorylation status of STING, TBK1, IRF3, and STAT1 at Y701. The STING agonist induced the phosphorylation of STING, TBK1, IRF3, and STAT1, most evident at 60 minutes post stimulation (**Fig. 4C**). At 60 minutes post diABZI stimulation total STAT1 levels fell. The ALPK1 agonist induced only a weak phosphorylation of TBK1 without affecting the phosphorylation status of the other proteins. Adding the ADP-heptose and diABZI together enhanced the early phosphorylation of TBK1 and increased the level of P-STAT1, while only marginally affecting the levels of P-STING and P-IRF3. The addition of ADP-heptose did not affect the diABZI triggered decline in STAT1 levels (**Fig. 4C**). These results indicate that ALPK1 activation does not directly activate the canonical STING pathway, but it can provide signals that amplify STING signaling.

**Fig. 4.**
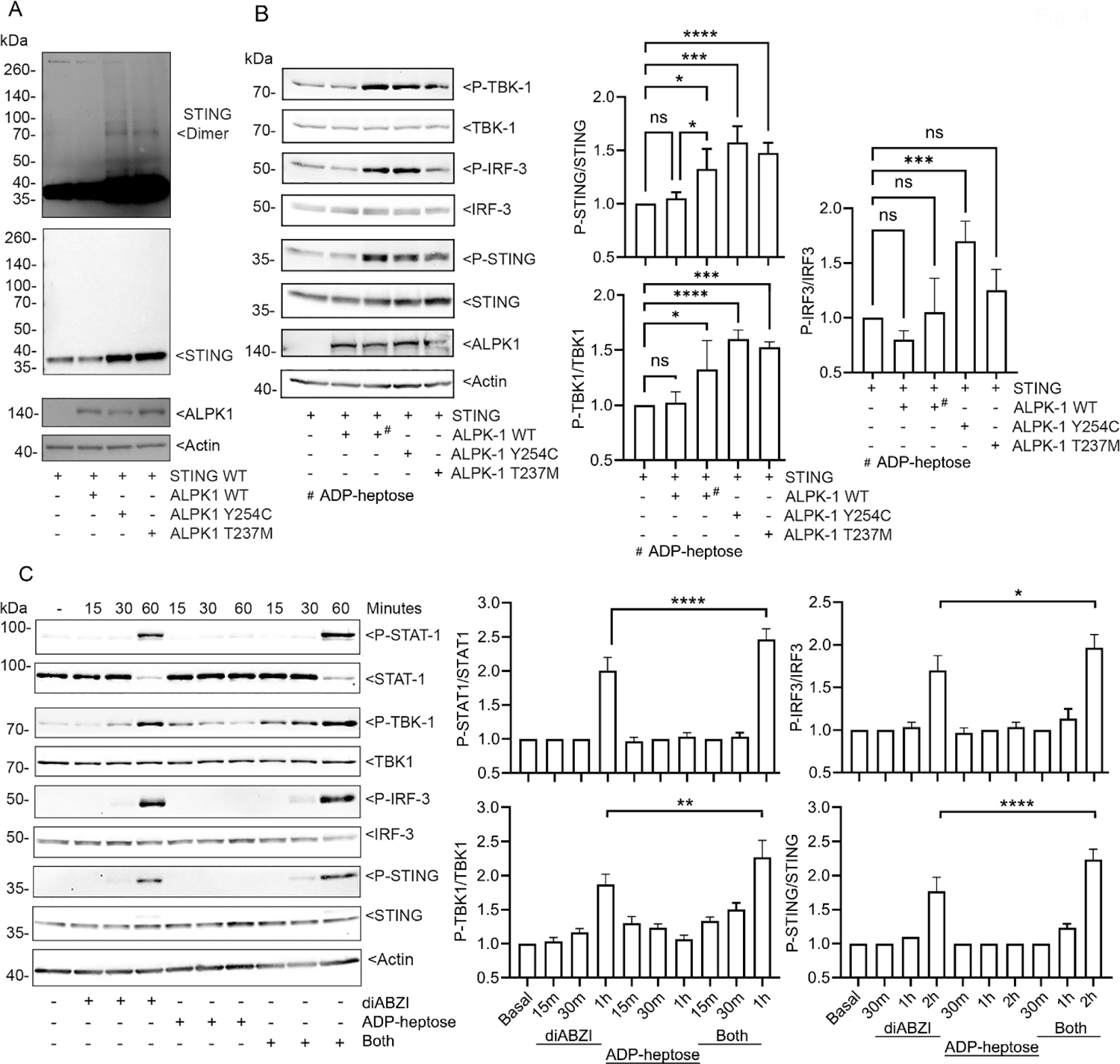
ALPK1 amplifies STING signaling. (**A-C**) Immunoblots and their quantification using cell lysates from HEK 293T cells or THP-1 cells transfected, or not, and treated with ADP-heptose and/or diABZI as indicated. Specific antibodies used to detect indicated proteins. (**A**) STING and the ALPK1 co-expression. Two exposures of STING immunoblot are shown to illustrate STING oligomers. One experiment of 3 biological replicates shown. (**B**) The indicated proteins were expressed in HEK 293T cells. P-STING, P-TBK1, P-IRF-3 the corresponding total proteins detected. ADP-heptose used at 1.5 µg/ml for 2h. One of three biological replicates shown. Quantification from all three. (**C**) PMA-differentiated THP-1 cells were stimulated with diABZI (1 µM), ADP-heptose (1.5 µg/ml) or both for 2 hrs. One of three biological replicates. Quantification from all three. Mean +/- S.E.M. ****, p<0.0001; ***, p<0.001; ***, p<0.05, NS-non-significant.

### ALPK1 enhances STING proton channel signaling and the STING-PERK (protein kinase regulated by RNA (PKR)-like kinase) pathway

STING signaling not only induces IRF3 and NF-κB dependent gene transcription, but it also causes proton leakage at the Golgi through a channel formed at the interface of the STING homodimer’s transmembrane domains (Liu et al., 2023; Xun et al., 2024). This channel activity is critical for LC3B lipidation, a post-translational modification important for autophagy, and monocyte NLRP3 inflammasome activation (Liu et al., 2023). Non lipidated LC3B, LC3B-I is cytosolic while the lipidated LC3B-II associates with autophagosomal membranes. Since LC3B-II runs slightly faster on SDS-PAGE, LC3B-I and -II can be distinguished by immunoblotting. Using HEK 293T cells we verified that diABZI-induced LC3B lipidation required STING (**Fig. 5A**). Treating the STING transfected cells with diABZI resulted in STING phosphorylation and an increase in LC3B-II. Co-expressing STING and ALPK1 Y254C or co-expressing STING with wild type ALPK1 and treating with ADP-heptose also enhanced LC3B-II levels (**Fig. 5B and C**). Compound 53 (C53) is a STING agonist that binds the putative channel interface blocking the STING-induced proton flux in the Golgi (Liu et al., 2023). Treating cells with C53 showed that the enhancement of LC3B-II by ligand-induced or mutation-induced ALPK1 activation depended upon STING proton channel activity (**Fig. 5D-F**).

**Fig. 5.**
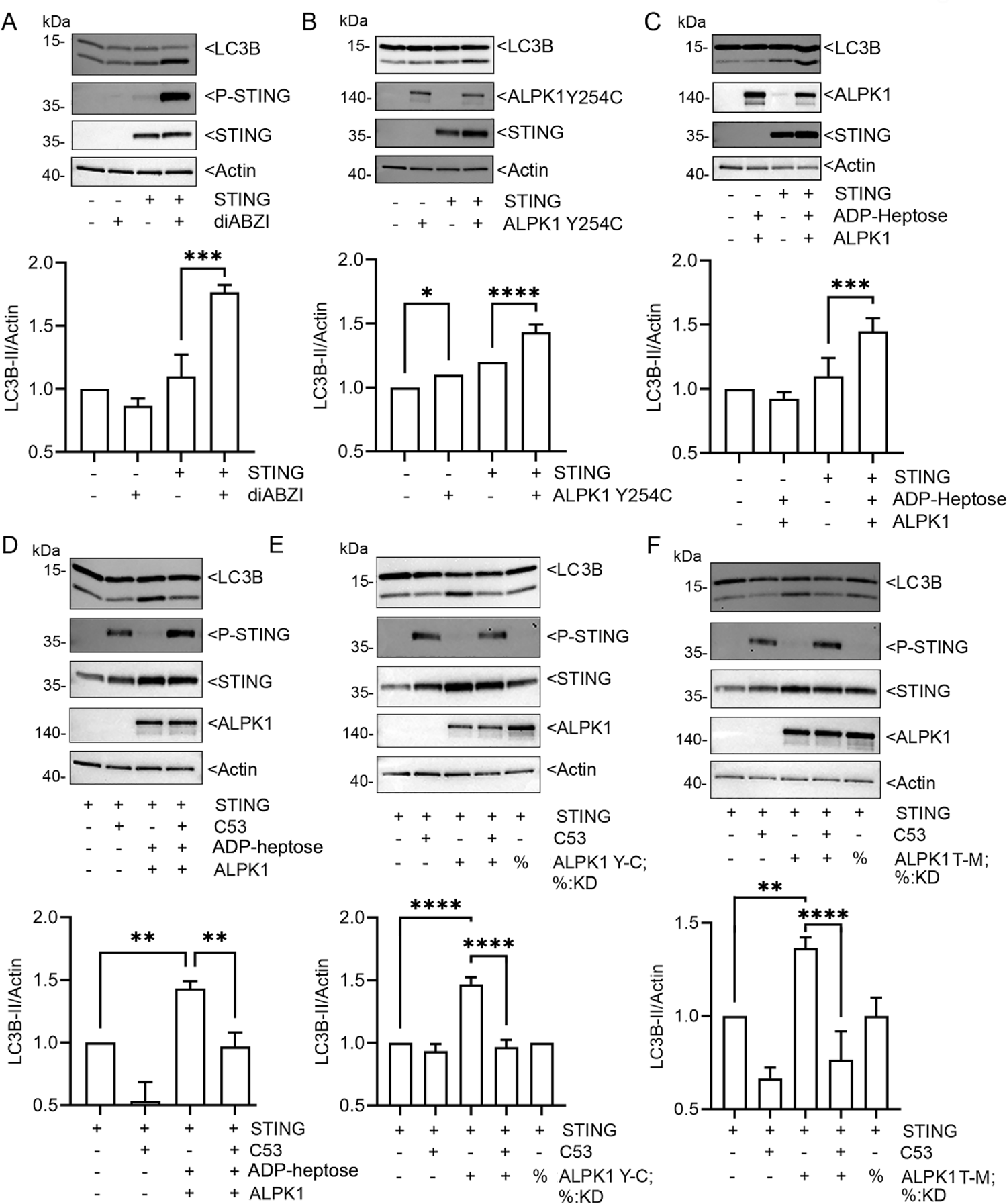
ALPK1 enhances STING induced LC3 lipidation. **(A-E)** Immunoblots and their quantification using cell lysates from HEK 293T cells transfected, or not, and treated as indicated. Specific antibodies used to detect indicated proteins. (**A-C**) HEK 293T cells transfected with STING, or not, and stimulated with diABZI (1 µM, 1 hour) or co-transfected with ALPK1 Y254C or ALPK1. The ALPK1 transfected cells stimulated with ADP-heptose (1.5 µg/ml, 1 hour). **(D-F**) HEK 293T cells transfected STING and either ALPK1, ALPK1 Y254C, or ALPK1 T237M or their corresponding kinase dead version. Some cells treated with C53 (10 µM) or ADP-heptose (1.5 µg/ml overnight). Each immunoblots from 1 of 3 biological replicates. Quantification from all three. Mean +/- S.E.M. ****, p<0.0001; ***, p<0.001; **, p<0.01; *, p>0.05, NS-non-significant.

Next, we used primed THP-1 cells to assess the role of STING proton channel activity and ALPK1 in NLRP3 inflammasome activation. The PMA priming enables stimuli to assemble NLRP3 inflammasomes. Following transfection of the ALPK1 gain of function mutants into the primed THP-1 cells we found an increase in IL-1β secretion. The addition of C53 or the NLRP3 inflammasome inhibitor MCC950 blocked those increases. These results indicate that activated ALPK1 triggers NLRP3 inflammasome activation via a STING-induced proton flux (**Fig. 6A**). Next, we stimulated THP-1 cells with diABZI, ADP-heptose or both agonists and measured IL-1β production. ADP-heptose alone had no effect, while diABZI slightly increased (1.4-fold) the IL-1β levels in the supernatant. Together the two agonists induced a 2.3-fold increase in IL-1β levels, which again depended upon STING proton channel activity (**Fig. 6B**).

**Fig. 6.**
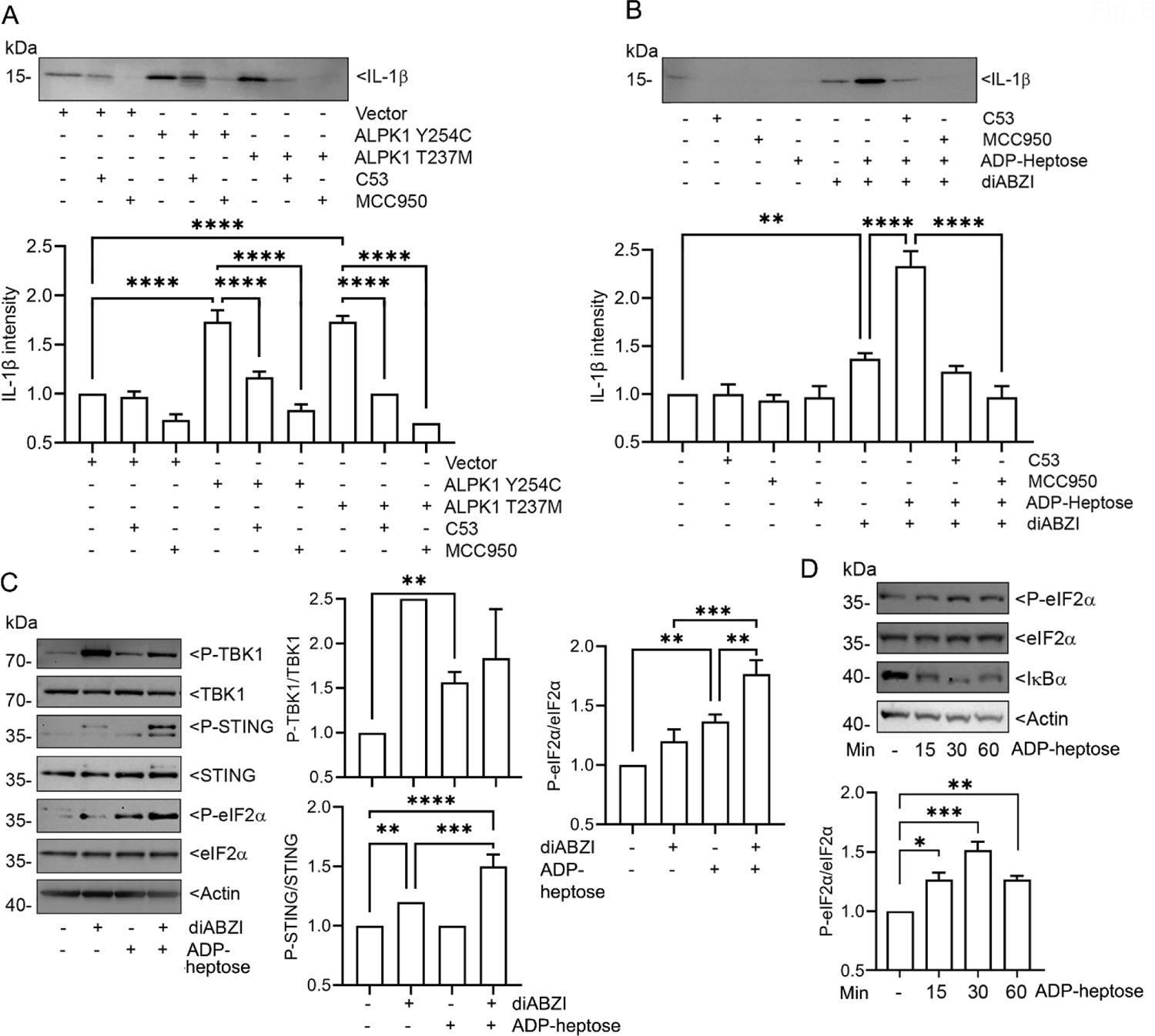
ALPK1 enhances STING-induced inflammasome activity and eIF2α phosphorylation. **(A-D)** Immunoblots and their quantification using cell lysates or supernatants from PMA-differentiated THP-1 cells, transfected or not, and treated as indicated. Proteins detected with specific antibodies. (**A**) Cells were transfected with gain function mutants of ALPK1 and treated, or not, with C53 (10 mM) or MCC9503 (5 mM) for 4 hours. Cell supernatants collected for IL-1β detection. (**B**) Cells treated with indicated combinations of C53 (10 mM), MCC9503 (5 mM), ADP-Heptose (1.5 µg/ml), or diABZI (1 mM) for 4 hours. Cell supernatants collected for IL-1β detection. (**C**) Cells stimulated with diABZI (1 mM), ADP-Heptose (1.5 mg/ml), or both for 1 hour. (**D**) Cells stimulated with ADP-heptose for indicated times. Immunoblots from 1 of 3 biological replicates. Quantitation from all three. Mean +/- S.E.M. ****, p<0.0001; ***, p<0.001; **, p<0.01; NS-non-significant.

Another non-canonical STING function is to activate the PERK-eIF2α (eukaryotic translation initiation factor 2α) pathway, which coordinates a translational program that promotes inflammatory and survival signaling. This occurs in the ER and is independent of the TBK1-IRF3 cascade (Zhang et al., 2022). To determine if ALPK1 signaling impacted this pathway we stimulated THP-1 cells with diABZI, ADP-heptose, or both and assessed eIF2α phosphorylation at serine 51 (S51). We found that ADP-heptose triggered eIF2α S51 phosphorylation even in the absence of diABZI and together they enhanced phosphorylation more so than did either agonist (**Fig. 6C**). To check whether the increase in eIF2α S51 phosphorylation depended upon STING we stimulated HEK 293T cells with ADP-heptose and assessed eIF2α S51 phosphorylation. We found a time dependent increase in eIF2α S51 phosphorylation that peaked at 30 minutes post stimulation (**Fig. 6D**). This argues that ALK1 mediated phosphorylation of eIF2α is independent of the STING/PERK/eIF2α pathway. Since ALPK1 is unlikely to directly phosphorylate eIF2α at S51, ALPK1 presumably engages an upstream signaling pathway that activates one of the known kinases that does.

### Further ALPK1 and STING signaling crosstalk

NF-κB activation prevents STING degradation by altering microtubule-mediated STING trafficking to lysosomes thereby enhancing STING signaling (Zhang et al., 2023). Conversely, STING agonist stimulate STING phosphorylation, which leads to STING degradation (Balka et al., 2023). To better understand how the ALPK1 and STING signaling pathways intersect we treated THP-1 cells with ADP-heptose, diABZI, or both and assessed P-TBK1, TBK1, P-STING, and STING levels at 2, 4, and 6 hours after their addition. The addition of ADP-heptose to diABZI minimally affected the kinetics of TBK1 phosphorylation or its expression level at 2, 4, or 6 hours after stimulation. However, both stimuli together increased STING phosphorylation and STING protein levels suggesting decreased STING degradation (**Fig. 7A**). The ALPK1 agonist alone increased STING protein amount (**Fig. 7A)**. Furthermore, ALPK1 protein levels rose following addition of the STING agonist despite the decline in STING protein levels (**Fig. 7B**).

**Fig. 7.**
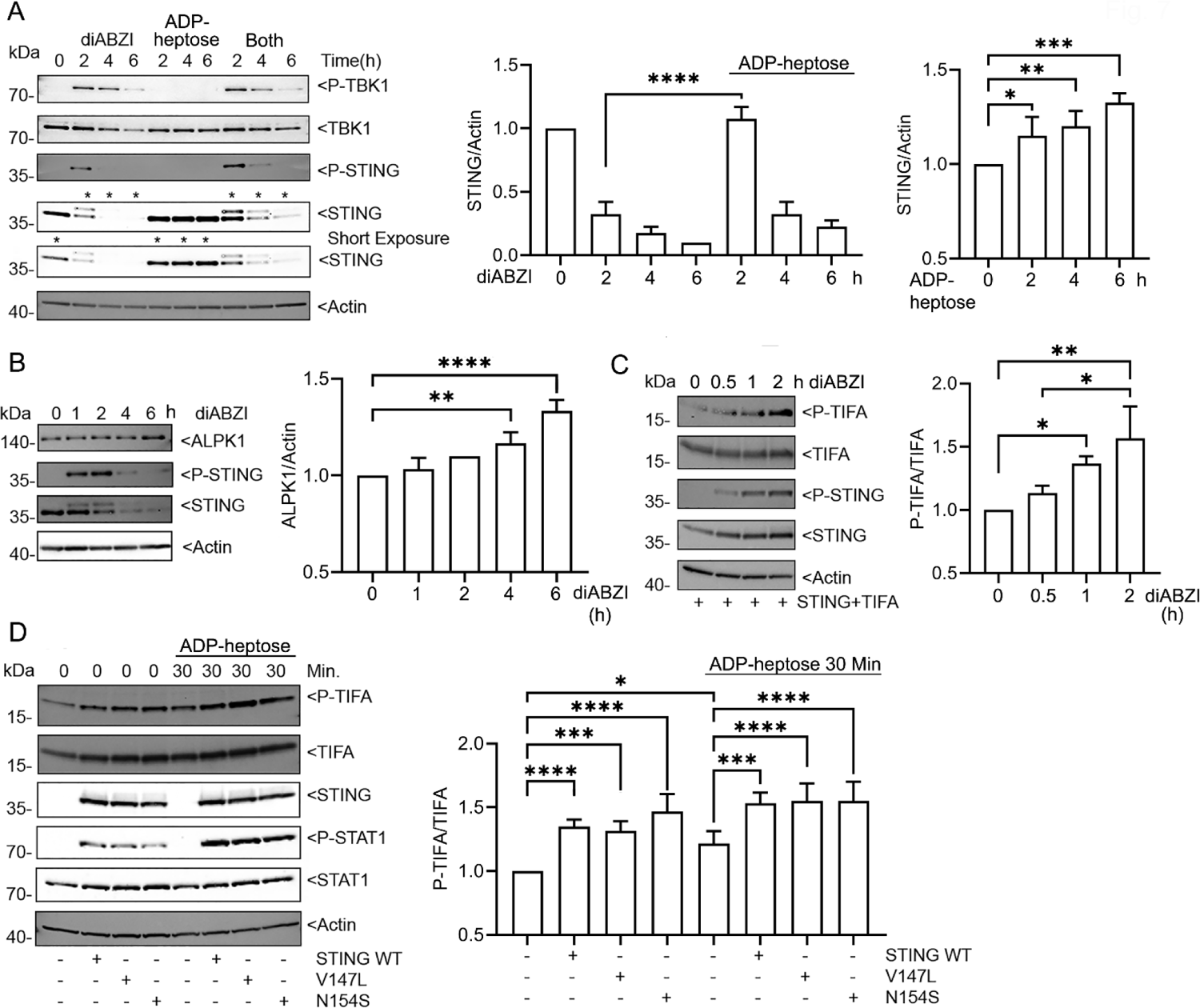
ALPK1 and STING crosstalk. (**A-D**) Immunoblots and their quantification using cell lysates from THP-1 or HEK 293T cells treated as indicated. Specific antibodies used to detect indicated proteins. (**A**) PMA-differentiated THP-1 cells treated for 2, 4, or 6 hours with diABZI (1 µM), ADP-heptose (1.5 µg/ml), or both. (**B**) PMA-differentiated THP-1 cells treated with diABZI for 1, 2, 4, or 6 hours. (**C**) HEK 293T cells transfected with STING and TIFA stimulated with diABZI (1 µM) or ADP-heptose (1.5 µg/ml) for 30 minutes, 1 hour, or 2 hours. (**D**) HEK 293T cells transfected with STING, STING V147L, or STING N154S and treated with ADP-heptose (1.5 µg/ml) for 30 minutes, or not. Each immunoblot from 1 of 3 biological replicates. Quantification from all three. Mean +/- S.E.M. ****, p<0.0001; ***, p<0.001; **, p<0.01; *, p>0.05, NS-non-significant.

TIFA is a downstream effector of ALPK1 as ALPK1 phosphorylates TIFA at threonine 9 (T9). Phosphorylated TIFA induces the oligomerization and polyubiquitination of TRAF6, which activates transforming growth factor-β-activated kinase 1 (TAK1) and the IκB kinase (IKK) complex, which phosphorylates IκBα. Phosphorylation of IκBα leads to its ubiquitination and proteasomal degradation which releases NF-κB to translocate to the nucleus (Garcia-Weber et al., 2023; Snelling et al., 2022; Zimmermann et al., 2017). STING signaling also activates the IKK complex similarly releasing NF-κB to translocate to the nucleus (Chen et al., 2016), however there is no known role for TIFA in STING signaling. To assess whether STING might also use TIFA to engage the NF-κB signaling pathway we checked the status of TIFA T9 phosphorylation following STING activation. We transfected HEK 293T cells with STING and TIFA, stimulated the cells with diABZI, and immunoblotted STING and TIFA along with their phosphorylated versions. At both 1 and 2 hours post stimulation TIFA phosphorylation at T9 as well as STING phosphorylation at S366 increased (**Fig. 7C**) suggesting that STING signaling had activated the TIFA signaling pathway. Next, we transfected STING wild-type and two gain-function mutants (Liu et al., 2014) into HEK 293T cells with and without ADP-heptose stimulation. The V147L mutation results in a gain-of-function change that increases the basal activity of STING even in the absence of its ligand and spontaneously activates downstream signaling pathways. The N154S mutation makes STING more responsive to activation. It can lead to constitutive activity or hypersensitivity to cGAMP or other agonists, amplifying downstream signaling. Expression of wild type STING and its mutants induced TIFA and STAT1 phosphorylation even in the absence of ADP-heptose. We noted little difference between the WT and the STING mutants (**Fig. 7D**). As expected, the addition of ADP-heptose in the absence of STING increased TIFA T9 phosphorylation, however, co-transfection of the STING constructs further increased the levels. The addition of ADP-heptose did not affect the STING enhanced phosphorylation of TIFA but did increase STAT1 phosphorylation (**Fig. 7D**). These data further support an interaction between the ALPK1/TIFA and the STING signaling pathway.

### ADP-Heptose and diABZI co-stimulation amplifies human monocyte cytokine production

To complement the cell line studies, we stimulated human peripheral blood monocytes with diABZI, ADP-heptose, or both ligands assessing early signaling by immunoblotting and cytokine production by multi-analyte flow cytometry. We immunoblotted monocytes from 3 normal individuals and have shown the results from 2 of them (**Fig. 8A**). As we observed with the THP-1 cells, diABZI plus ADP-heptose increased the levels of P-TBK1, P-IRF3, and P-STING compared to diABZI alone. The ADP-heptose treatment alone increased STING protein expression. In contrast to the THP-1 cells, ADP-heptose alone increased P-STAT1 level and combining with it with diABZI reduced the ADP-heptose mediated increase. Next, we purified monocytes from the peripheral blood of 5 human donors. Shown are results from one donor whose monocytes were cultured in triplicate, and each culture assayed in triplicate. Among the various cytokines we tested only IL-10 and CXCL8 levels rose in the ADP-heptose alone treated monocyte cell supernatants (**Fig 8B**). This was the case for all the tested individuals **(supplementary fig. S1**). Together ADP-Heptose and diABZI significantly increased all the cytokines we tested compared to either alone (**Fig. 8B**). Due to variability in responsiveness, the combined data from the 5 donors showed a significant synergy between ADP-heptose and diABZI in the production of IL-1β, IL-6, TNF-α CXCL8, interferon-λ1, interferon-β, interferon-γ, and GM-CSF (**supplementary Fig. S1**). Overall, the human monocyte studies support the cell line data previously detailed.

**Fig. 8.**
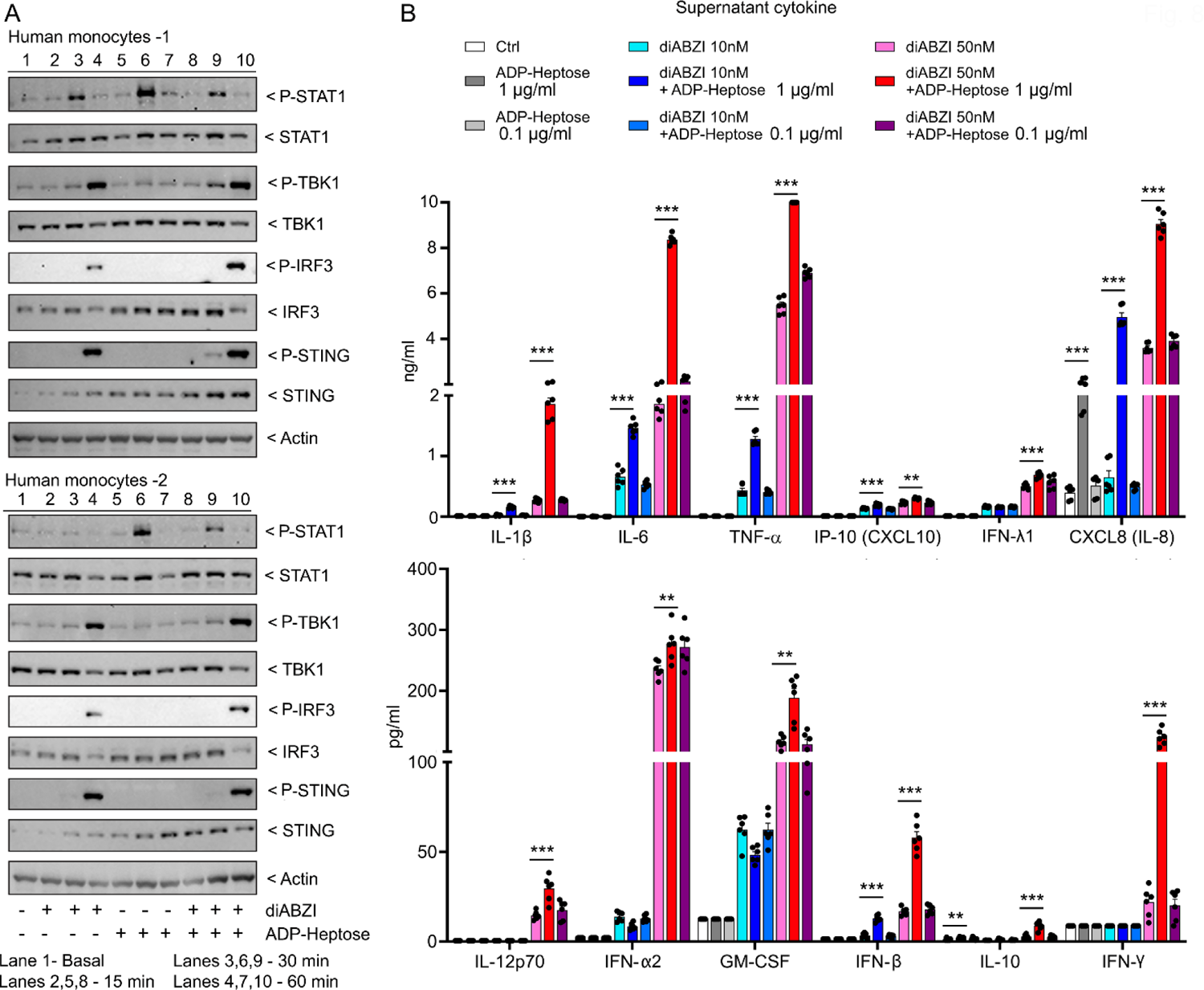
ADP-heptose amplifies diABZI induced cytokine production by human monocytes. **(A-B**) Human monocytes purified from peripheral blood stimulated with ADP-heptose, diABZI, or both. Cell lysates immunoblotted or 24-hour conditioned cell supernatants collected for cytokine production. (**A**) Cell lysates prepared from 2 donors stimulated for various duration and immunoblotted for indicated proteins. (**B**) Three cell supernatants from 1 donor each assayed in triplicate for indicated cytokines by multi-analyte flow cytometry. Mean +/- S.E.M. ***, p<0.001; **, p<0.01; *, p>0.05, NS-non-significant.

## Discussion

This study reveals a previously unrecognized, bidirectional crosstalk between ALPK1 and STING, two cytosolic immune sensors traditionally associated with distinct innate immune pathways. To our knowledge, this is the first demonstration of reciprocal amplification between ALPK1-mediated bacterial metabolite sensing and STING-mediated DNA sensing pathways in human disease. By leveraging mechanistic cellular models alongside clinical phenotyping of ROSAH syndrome, a monogenic autoinflammatory disorder driven by activating ALPK1 mutations, we establish that ALPK1 activation amplifies STING signaling, while STING activation reciprocally modulates ALPK1/TIFA/NF-κB signaling.

Previous studies indicate that canonical NF-κB activation can inhibit STING trafficking to lysosomes for degradation via microtubule depolymerization, resulting in sustained STING activity and a heightened interferon response (Zhang et al., 2023). Consistent with these observations, we found that ALPK1 activation enhanced both canonical and non-canonical STING signaling. ALPK1 gain-of-function mutants activated an IFN-β reporter in a STING-, ALPK1 kinase-, and NF-κB p65-dependent manner. Moreover, these mutants potentiated STING-agonist-induced phosphorylation of STING, TBK1 and IRF3, highlighting a robust amplification of canonical STING responses.

Beyond type I interferon signaling, ALPK1 also modulated STING’s noncanonical outputs. STING-driven LC3B lipidation and inflammasome activation require its translocation to the Golgi and channel activity, but not on TBK1 or IRF3 (Liu et al., 2023). While NF-κB helps prime monocyte and macrophages for inflammasome activation, it has no known direct role in LC3B lipidation. ALPK1 gain-of-function mutants and agonist activated ALPK1 increased LC3B lipidation in a manner dependent on STING, its channel function (as inhibited by C53), and ALPK1 kinase activity. Whether this effect is mediated through TIFA/NF-κB or another downstream ALPK1 pathway remains to be determined.

While STING channel activity mediates both LC3B lipidation and NLRP3 inflammasome activation, these pathways diverge downstream of the proton leakage. The precise mechanism by which STING-induced proton flux activates NLRP3 remains unclear. Following STING’s translocation to the Golgi, cytosolic NLRP3 also translocates to the Golgi where it initiates inflammasome assembly - a process blocked by the STING channel inhibitor C53, implicating proton leakage as the inciting event. Expression of gain-of-function ALPK1 mutants in primed THP-1 cells enhanced IL-1β production, an effect that required STING channel activity. Although ADP-heptose alone did not increase IL-1β section in these cells, adding a suboptimal amount of diABZI to the ADP-heptose did, and either C53 or the NLRP3 antagonist blocked the increase. Since ALPK1 activation augments both STING mediated inflammasome activation and LC3B lipidation, ALPK1 signaling may directly target STING channel activity. Interestingly, the influenza virus M2 protein—another proton channel—also induces LC3B lipidation and NLRP3 activation (Beale et al., 2014; Ichinohe et al., 2010). Whether ALPK1 activation can modulate the activity of such channels is an intriguing area for future study, especially given the potential for bacterial metabolite sensing to exacerbate STING-mediated inflammation during viral infections like influenza.

While most STING signaling occurs after it translocates to the Golgi, the initial binding of cGAMP to ER-localized STING directly activates PERK, one of four known eIF2α kinases. This precedes TBK1-IRF3 activation and does not involve the unfolded protein response, a potent PERK activator. Once activated, PERK phosphorylates eIF2α, thereby activating a translation program that favors inflammation and cell survival and links the innate immune responses to broader stress-adaptive pathways (Zhang et al., 2022). In primed THP-1 cells, we found that both ADP-heptose and diABZI increased eIF2α phosphorylation, with further enhancement when used together. ADP-heptose also induced this phosphorylation in STING-deficient HEK293T cells, suggesting that ALPK1 activates a separate signaling pathway that converges on eIF2α. One strong candidate is the heme-regulated inhibitor (HRI), another eIF2α kinase involved in pro-inflammatory responses to intracellular bacteria, acting downstream of MAVS and TRIF but not STING or MyD88 (Abdel-Nour et al., 2019). Current studies are investigating how ALPK1 engages this signaling arm and whether dysregulated eIF2α phosphorylation contributes to immune dysfunction or chronic inflammation in individuals with ALPK1 mutations.

Our data also reveal reciprocal regulatory effects. ADP-heptose increased endogenous STING levels and delayed its degradation, likely through NF-κB-mediated augmentation of STING transcription and inhibition of lysosomal trafficking (Chen et al., 2023; Zhang et al., 2023). Similarly, STING agonist treatment of THP-1 cells increased the amount of endogenous ALPK1, potentially via STING/NF-κB pathway, which triggers IL-6 and IL-6 receptor expression and STAT3 activation (Hong et al., 2022). STAT3 in-turn can bind the promoter of ALPK1 increasing transcription. Finally, exposure to STING agonists induced phosphorylation of TIFA at Thr9, suggesting that STING-driven signaling may augment TIFAsome formation—a scaffolding event known to enhance NLRP3 inflammasome responses (Lin et al., 2016). These reciprocal influences point to a tightly interconnected signaling circuit capable of finely tuning innate immune activation.

While PRR crosstalk such as that observed between ALPK1 and STING likely provides critical advantages during acute infections, chronic or dysregulated activation may carry significant long-term risks, fueling chronic inflammation, tissue damage, or degeneration. The ALPK1–STING signaling axis thus emerges not only as a powerful innate immune amplifier but also as a potential driver of disease across a spectrum of inflammatory and degenerative conditions.

This signaling circuit may be particularly relevant in immune-privileged tissues such as the CNS, where tolerance to chronic inflammation may be more limited. ROSAH syndrome, characterized by constitutive ALPK1 activation, offers a unique window into the in vivo consequences of this signaling axis. In our expanded ROSAH cohort, most individuals exhibited accelerated intracranial mineralization—a process that typically occurs with aging but can be markedly accelerated in type I interferonopathies such as Aicardi-Goutières syndrome and in congenital infection with viruses like CMV (Alarcon et al., 2013; La Piana et al., 2016). Notably, all ROSAH patients who underwent lumbar puncture had elevated CSF neopterin, a marker of IFN-γ–mediated activation of macrophages and astrocytes (Dale et al., 2009; Han et al., 2022; Martin et al., 2025), despite the absence of a strong interferon signature in peripheral blood. This divergence between systemic and CNS immune markers highlights the growing recognition of tissue-specific immune responses, suggesting that chronic ALPK1–STING crosstalk may exert especially pronounced effects in immune-restricted compartments like the CNS.

Our findings also carry important implications for diseases beyond rare monogenic conditions. Dysregulated STING signaling impacts a wide range of cellular processes—including autophagy, metabolic regulation, and DNA damage repair—and has been implicated in infectious, autoimmune, malignant, fibrotic, and neurodegenerative disorders (Chen and Xu, 2023; Jeltema et al., 2023). Yet, the upstream triggers that drive STING activation in many of these contexts remain incompletely understood. Our data suggest that crosstalk between ALPK1 and STING may be particularly relevant in the central nervous system, a finding of interest given that STING activation has been shown to promote aging-related inflammation and neurodegeneration (Gulen et al., 2023). In parallel, recent studies have shown that aging is associated with increased gut permeability and systemic dissemination of microbial metabolites including ADP-heptose (Agarwal et al., 2025). While existing work has linked increasing in circulating ADP-heptose to age-associated clonal hematopoiesis, our findings raise the possibility that these metabolites may also activate the ALPK1–STING axis in the CNS and other tissues, thereby contributing to additional age-associated inflammatory and degenerative processes.

In conclusion, we demonstrate a previously unrecognized bidirectional crosstalk between ALPK1 and STING that enhances canonical and non-canonical innate immune pathways. While initially elucidated in ROSAH syndrome, these insights hold broader implications for understanding and potentially treating a range of inflammatory and degenerative diseases. Our findings provide a foundation for future therapeutic strategies that selectively target this signaling axis to modulate immune responses in a tissue-specific and context-dependent manner.

## Materials and Methods

### Patients, clinical and laboratory evaluation

In this retrospective cohort study, we collected data on patients with ROSAH enrolled on the NIH clinical trial number: NCT00001373. Data capture focused on demographics and MRI brain scan findings. Previously banked patient plasma was used for CXCL10 measurement. As previously described, the nanostring platform was used to calculate a 28-gene IFN score from whole blood. (Kim et al., 2018). Additionally, five individuals with ROSAH and headaches underwent lumbar puncture for CSF evaluation.

### Human monocytes isolation

Peripheral blood mononuclear cells(PBMCs) were isolated by Ficoll-Hypaque density centrifugation from buffy coats or whole blood donated by healthy adult donors. Buffy coats and whole blood were provided by the NIH Department of Transfusion Medicine (DTM)-approved protocol (Institutional Review Board of the NIAID). Monocyte cell population were obtained by negative selection (>97% purity, Stem Cell Technologies).

### Reagents

ADP-Heptose (InvivoGen) was used at the indicated concentrations. diABZI (Selleckchem) was used at 1 μM, C53 (Cayman chemical) at 10 µM, and MCC950 (Selleckchem) at 5 µM. Primary Abs used for immunoblot experiments were directed against STING (#13647; 1:2000), Phospho-STING S366 (#50907; 1:1500), TBK1 (#38066; 1:2000), Phospho-TBK1 S172 (#5483; 1:1500), IRF-3 (#4302; 1:1000), Phospho-IRF-3 S386 (#37829; 1:1000), STAT1 (#14994; 1:1500), Phospho-STAT1 Y701(#9167; 1:1500), eIF2α(#5324; 1:1500), Phospho-eIF2α S51(#3398; 1:1000), IL-1β (#12703; 1:1000), and LC3B (#43566; 1:1000). The above Abs were purchased from Cell Signaling Technology. The antibodies that recognize ALPK1 were purchased from Abcam (ab236626; 1:5000). An anti-rabbit IgG HRP-linked antibody (Cell Signaling Technology; #7074; 1:10000) was used for immunoblots. HEK 293T and THP-1 cells were obtained from the American Type Culture Collection. HEK 293T cells were maintained in DMEM supplemented with 10% FBS (Invitrogen). THP-1 cells were maintained in RPMI-1640 with 10% FBS (Invitrogen) added Sodium Pyruvate 1mM (Corning) and 2-Mercaptoethanol 2 µl (Sigma). PMA 10 nM was used to treat THP-1 cells 2h for macrophage differentiation. After which the cells were used for transfection experiments and/or agonist stimulation. The IFN-β luciferase reporter plasmid was a gift from Dr. Saori Suzuki (University of Tokyo, Tokyo, Japan). The NF-κB luciferase reporter was provided by Dr. Ulrich Siebenlist (National Institutes of Health). STING plasmid was a gift from Dr. Raphaela Goldbach-Mansky. The plasmids were transiently transfected into the cells using X-tremeGENE HP DNA Transfection Reagent (Roche) following the manufacturer’s protocol.

### Human monocytes treatment and quantification of cytokines

Primary blood monocytes were cultured in RPMI-1640 medium supplemented with 10% (v/v) heat inactivated fetal calf serum (HI FCS, ThermoFisher), 1% (v/v) Penicillin-Streptomycin (P/S, ThermoFisher) and 2 mM L-glutamine (ThermoFisher). Monocytes at 200,000 cells per well (1x 10^6^ cells/ml) were plated in 96 well plates and stimulated with diABZI (10nM, or 50nM, SelleckChem) for 24 h either in the presence or absence of ADP-heptose (0.01 µg/ml, or 1 µg/ml, InvivoGen). Culture supernatants were collected, and non-adherent cells removed by centrifugation. Samples were stored at −80 °C until analysis. The cultured supernatants were collected, and cytokine levels determined using the bead-based immunoassays LEGENDplex Human Anti-Virus. Response Panel (BioLegend). The assays were performed in 96-well plates following the manufacturer’s instructions. For measurements, a FACSCelesta SORP flow cytometer (BD Biosciences) was employed, and data were evaluated with the LEGENDplex Data Analysis software (BioLegend).

### p65 Crispr knock out (P65-/-) HEK 293T cell line generation

A pSpCas9(BB)-2A-GFP (PX458) from Addgene (#48138) was used to insert p65 targeted gRNA (AGCGCCCCTCGCACTTGTAG), then the construct was transfected into HEK 293T cells for 48h. The GFP positive cells were sorted and seeded into 96 well plate for culture. The p65^-/-^ single clone cells were verified by anti-p65 antibody immununoblotting.

### Luciferase reporter assay

HEK 293T cells (250 cells/well) were seeded in 48-well plates. The following day, the cells were co-transfected with 2 ng luciferase reporter plasmid, 1 ng Renilla luciferase internal control vector phRL-TK (Promega), and each of the indicated plasmids with X-tremeGENE HP DNA Transfection Reagent (Roche). Empty vector pcDNA3 was used to maintain equivalent amounts of DNA in each well. The reporter gene assay was performed 24 h after transfection and analyzed by a dual-luciferase reporter assay on the Mithras LB 840 Multimode Microplate Reader (Berthold). Each experiment was replicated a minimum of six times.

### Immunoblot analysis

For standard immunoblotting, the cells were lysed in a buffer of 20 mM HEPES (pH 7.4), 50 mM β-glycerophosphate, 1 mM Na3VO4, 0.5% (v/v) Triton X-100, 0.5% (v/v) CHAPS, and 10% (v/v) glycerol with a protease inhibitor mixture tablet. For the LC3 immunoblot analysis, the cells were lysed with the same buffer plus 0.25% SDS. The lysates were separated by SDS-PAGE and transferred to nitrocellulose membrane by iBLOT Gel Transfer System (Invitrogen). The membrane was incubated with 5% nonfat milk w/v in TBS buffer (25 mM Tris-HCl [pH 7.5], 150 mM NaCl, 0.1% Tween 20) for 1 h, and then reacted with the primary Ab in TBS buffer with 2.5% nonfat milk or 5% BSA w/v for overnight by shaking at 4°C. The appropriated second Abs conjugated to HRP was used to detect the protein of interest via ECL on iBRIGHT FL1000. Quantification of band intensity was done using ImageJ.

### Imaging

Hela cells or HEK293T cells were seeded in an 8-chambered coverglass system (Cellvis, C8-1.5H-N) overnight. Cells were transfected with ALPK1-Emerald or ALPK1 Y254C-Emerald, and Sting-mRuby plasmids at 0.32 μg/mL and 0.5 μg/mL, respectively, using X-treme GENE HP DNA Transfection Reagent (Roche). Cells were imaged after approximately 40 hours of protein expression. For live-cell studies, CO2 levels were maintained at 5%, and the temperature was kept at 37°C. Imaging was performed using a Leica SP8 inverted 5-channel confocal microscope equipped with ultra-sensitive hybrid detectors (Leica Microsystems, Buffalo Grove, IL), 488nm and 561nm lasers, and a 63x oil immersion objective (NA 1.4). Images of dynamic cell interactions were recorded with four z-stacks (3D) with ∼1 micron step interval over time for 1-2 hours (4D). Adaptive Focus Control allowed for the addition of reagent without disrupting focus.

### Statistical analysis

Prism (GraphPad Prism9) was used for all statistical analyses. All experiments were repeated a minimum of three times.

## Acknowledgements

We thank Clinton Bradfield (LISB,NIAID, NIH) as for construction of the Emerald tagged ALPK1 and Margery Smelkinson (RTB, NIAID, NIH) for assistance with the confocal microscopy.

## Author Contribution

CSS designed most of the experiments and performed all the immunoblots and assisted in writing the manuscript; CK managed the patients and assisted with the experimental design and helped with manuscript writing; NNH performed all the imaging experiments; IYH performed all the cytokine assays; DAH analyzed the MRIs; DLK provided patients, reviewed the study, and provided expert advice; and JHK wrote the manuscript and oversaw the project.

## Resource and Material Availability

Further information and requests for resources and reagents should be directed to and will be fulfilled by the Lead Contact, John H. Kehrl.

