## Supplementary material for "Crosstalk Between ALPK1 and STING: A Synergistic Axis in Innate Immune Activation and Human Inflammatory Disease": Figure and video captions

**Contents**

Supplement fig. 1, parts 1-3.

**Movie legends**

Supplement movie 1-4.

### Supplement fig. 1

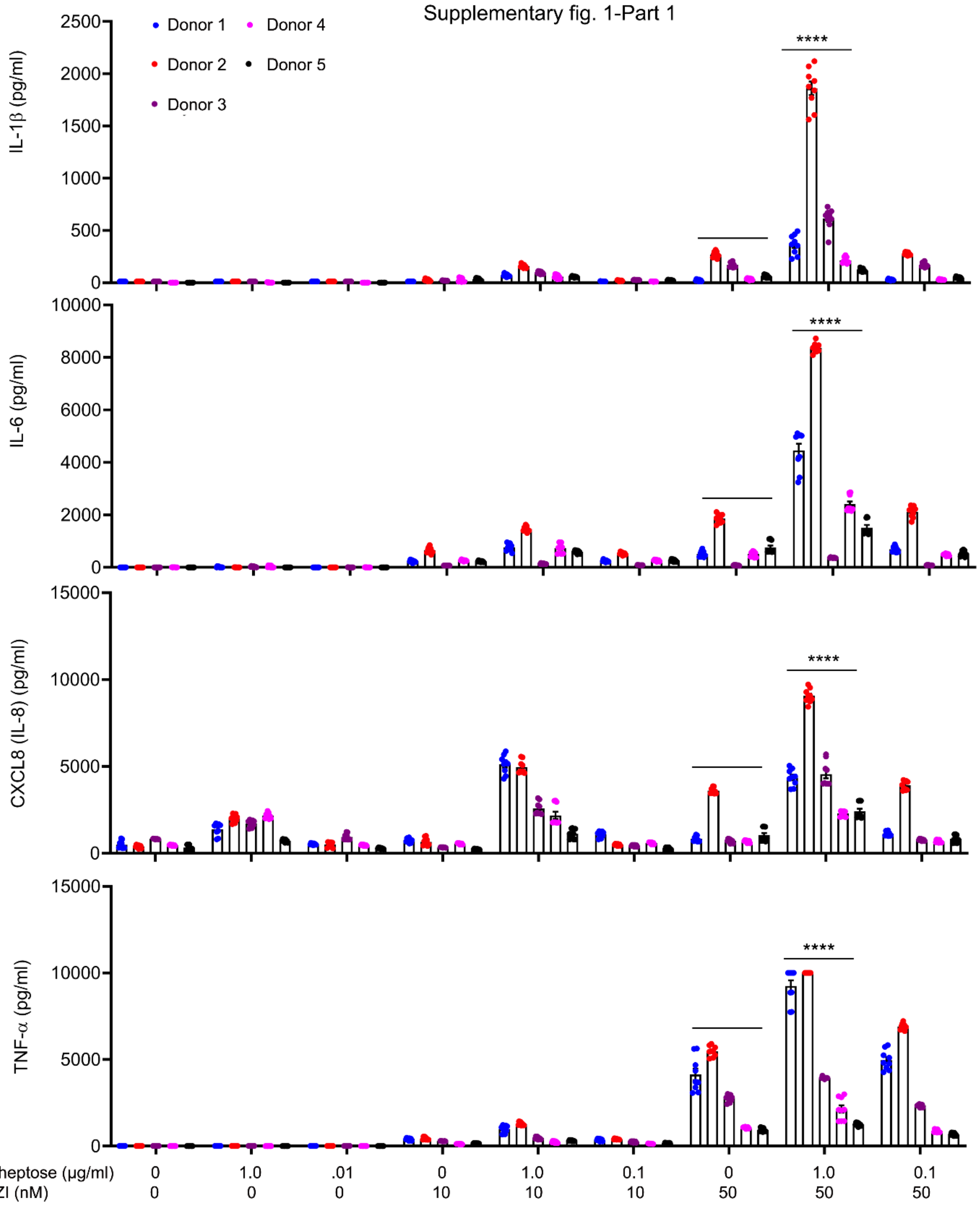

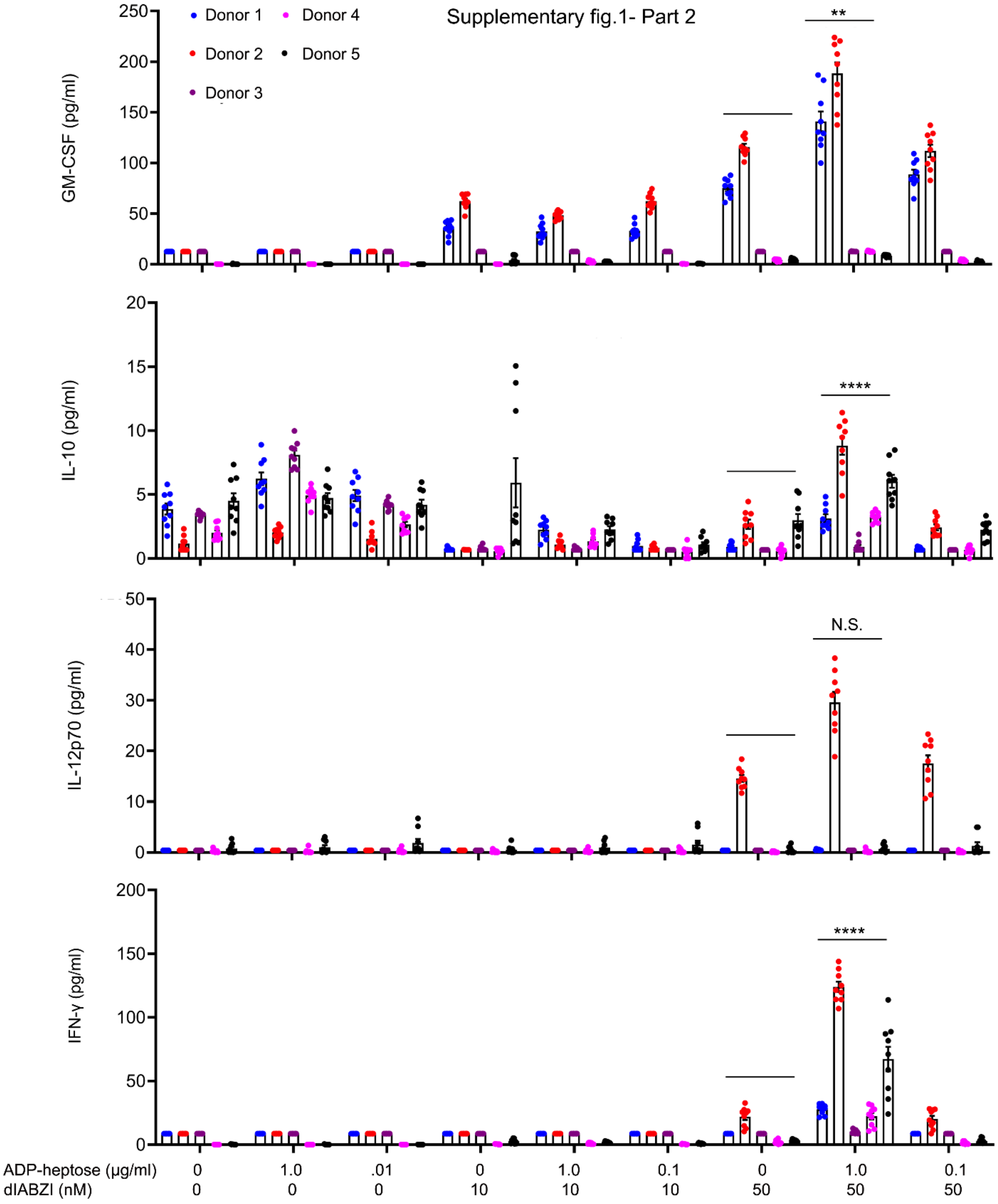

Supplementary fig.1- Part 2
Donor 1
Donor 2
Donor 3
Donor 4
Donor 5
GM-CSF (pg/ml)
**
IL-10 (pg/ml)
****
IL-12p70 (pg/ml)
N.S.
IFN-γ (pg/ml)
****
ADP-heptose (μg/ml) 0 1.0 .01 0 1.0 0.1 0 1.0 0.1
dIABZI (nM) 0 0 0 10 10 10 50 50 50

Supplementary fig. 1-Part 3

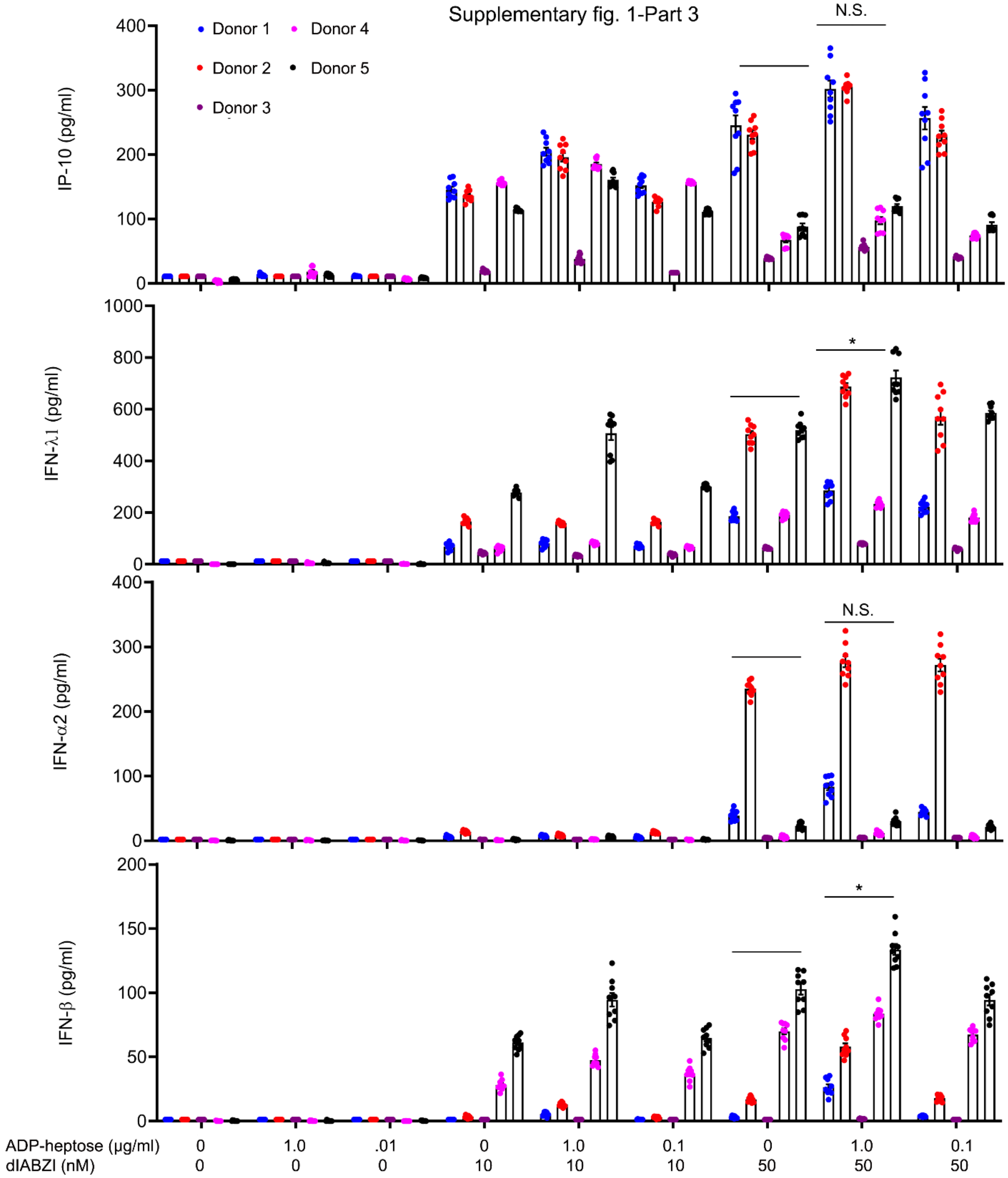

**Supplementary fig. 1. Cytokine production by human monocytes from 5 donors.**

Human monocytes purified from peripheral blood stimulated with ADP-heptose, diABZI, or both. Three cell supernatants from each donor each assayed in triplicate for indicated cytokines by multi-analyte flow cytometry. Statistics for comparison between diABZI (50 nM) and diABZI (50 nM) plus ADP-heptose (1.0 µg/ml) using data from all 5 donors.

**Movie 1**. ALPK1 and STING co-expression in HeLa cells. Live cell confocal imaging of HeLa cells transiently transfected with ALPK1-emerald (green) and STING-mRuby (pink) was performed. On day 2 following transfection the cells were imaged every 2 minutes for 2 hours. ADP-heptose 1.5 µg/ml was added at frame 10. Movie frame rate is 10 frames/second. Scale bar 10 microns. A single confocal image is shown for each frame.

**Movie 2**. ALPK1 Enhances STING expression in HeLa cells. Same live cell imaging as shown in movie 1, but the expression level of STING was mapped using the color table file Custom 1 in the Imaris program. The color gradient is shown in Figure 3. Imaging and frame rate the same as movie 1.

**Movie 3**. ALPK1 and STING Co-expression in HeLa cells. Live cell confocal imaging of HeLa cells transiently transfected with ALPK1-emerald (green) and STING-mRuby (pink) was performed. On day 2 following transfection the cells were imaged every minute for 90 minutes, movie spans the last 55 minutes. ADP-heptose 1.5 µg/ml was added just prior to initiating imaging. Movie frame rate is 10 frames/second. Scale bar 2 microns. A single confocal image is shown for each frame. Green-pink overlap is white.

**Movie 4**. ALPK Y254C and STING co-expressed in HEK293T cells. Live cell confocal imaging of HEK293T cells transfected 2 days previously with ALPK1 Y254C-Emerald and STING-mRuby and treated with diABZI (1 µM) at time point 18. Cells were imaged every minute for 90 minutes. Movie frame rate is 10 frames/second. A single confocal image is shown for each frame. Scale bar 4 microns. Green-pink overlap is white.
